# Distinct energy-coupling factor transporter subunits enable flavin acquisition and extracytosolic trafficking for extracellular electron transfer in *Listeria monocytogenes*

**DOI:** 10.1101/2022.11.04.515267

**Authors:** Rafael Rivera-Lugo, Shuo Huang, Frank Lee, Raphaël Méheust, Anthony T. Iavarone, Ashley M. Sidebottom, Eric Oldfield, Daniel A. Portnoy, Samuel H. Light

## Abstract

A variety of electron transfer mechanisms link bacterial cytosolic electron pools with functionally diverse redox activities in the cell envelope and extracellular space. In *Listeria monocytogenes*, the ApbE-like enzyme FmnB catalyzes extracytosolic protein flavinylation, covalently linking a flavin cofactor to proteins that transfer electrons to extracellular acceptors. *L. monocytogenes* uses an energy-coupling factor (ECF) transporter complex that contains distinct substrate-binding, transmembrane, ATPase A, and ATPase A’ subunits (RibU, EcfT, EcfA, and EcfA’) to import environmental flavins, but the basis of extracytosolic flavin trafficking for FmnB flavinylation remains poorly defined. In this study, we show that the proteins EetB and FmnA are related to ECF transporter substrate-binding and transmembrane subunits, respectively, and are essential for exporting flavins from the cytosol for flavinylation. Comparisons of the flavin import versus export capabilities of *L. monocytogenes* strains lacking different ECF transporter subunits demonstrates a strict directionality of substrate-binding subunit transport but partial functional redundancy of transmembrane and ATPase subunits. Based on these results, we propose that ECF transporter complexes with different subunit compositions execute directional flavin import/export through a broadly conserved mechanism. Finally, we present genomic context analyses which show that related ECF exporter genes are distributed across the Firmicutes phylum and frequently co-localize with genes encoding flavinylated extracytosolic proteins. These findings clarify the basis of ECF transporter export and extracytosolic flavin cofactor trafficking in Firmicutes.

## Background

Flavins are an important family of redox-active cofactors that catalyze electron transfer in diverse enzymes.^1^ Flavin mononucleotide (FMN) and flavin adenine dinucleotide (FAD) are synthesized from the precursor riboflavin (also known as vitamin B_2_). FMN and FAD are the most common type of protein-bound flavins and are nearly ubiquitous throughout the three domains of life. While the role of flavin-binding proteins (flavoproteins) in cytosolic redox activities is well established, the importance of flavins for extracytosolic activities in prokaryotic biology has become increasingly apparent.^2–4^

In prokaryotes, many extracytosolic flavoproteins are post-translationally linked to their flavin cofactors (flavinylated) through the action of the enzyme ApbE.^5^ ApbE specifically uses FAD as a substrate, catalyzing a reaction that links the FMN portion of the molecule to a serine/threonine residue in substrate proteins via a phosphodiester bond (**Figure 1A**).^6^ Approximately 50% of sequenced bacterial genomes encode proteins flavinylated by ApbE, with ApbE substrates having been implicated in a wide array of redox-dependent activities.^4^ For example, *Rhodobacter* nitrogen fixation (Rnf) and NADH:quinone oxidoreductase (Nqr) are prominent multi-subunit complexes with ApbE-flavinylated subunits that possess important roles in diverse bacteria and energy metabolisms.^7,8^

**Figure 1.**
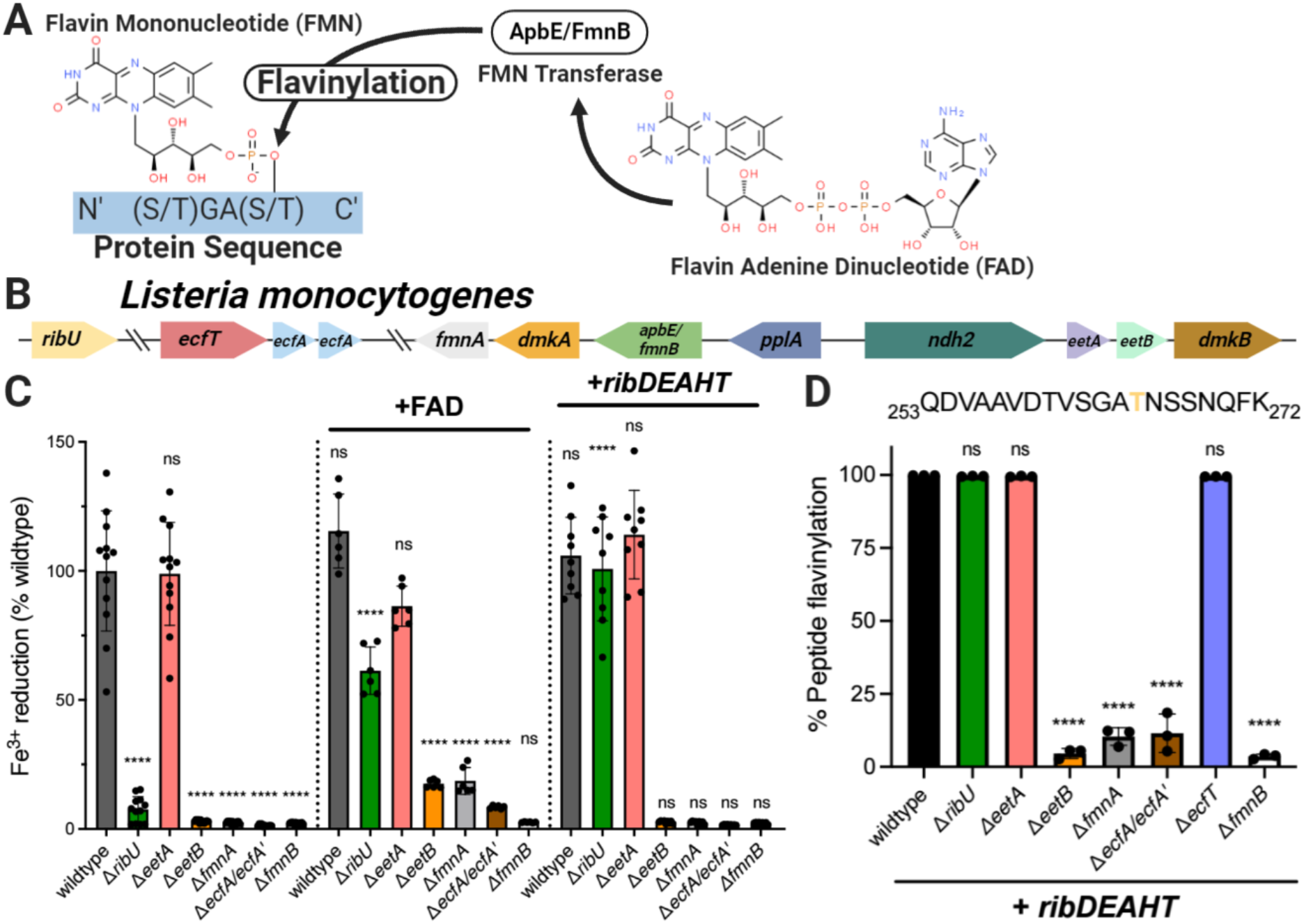
ECF transporter subunits and *eetB* are essential for provisioning flavins to the extracytoslic ApbE homolog FmnB in *L. monocytogenes*. (A) Reaction catalyzed by AbpE/FmnB flavin transferases. (B) Genomic organization of *L. monocytogenes* genes addressed in this study. (C) Ferric iron reductase activity of *L. monocytogenes* strains grown in chemically defined media. The dotted lines separate flavin auxotroph (wildtype) strains (*n*=4), flavin auxotroph strains supplemented with extracellular FAD (*n*=2), and flavin prototroph (+*ribDEAHT*) strains (*n*=3). For the non-complemented strains, statistical significance was determined using one-way ANOVA and Dunnett’s posttest using wildtype as the control. For the FAD and *ribDEAHT* complemented conditions, statistical significance was determined by performing a *t*-test to the parental non-complemented condition. (D) PplA flavinylation levels in flavin prototroph (+*ribDEAHT*) *L. monocytogenes* strains as assessed by MS. The data show the means and SDs of three independent experiments. Statistical significance was determined using one-way ANOVA and Dunnett’s posttest using wildtype as the control. ****P < 0.0001; ns, not significant, P > 0.05.

Extracellular electron transfer (EET) describes a class of microbial activities that result in the transfer of electrons from the cytosol to the outside of the cell and which often function in anaerobic respiration. We previously found that the foodborne pathogen *Listeria monocytogenes* possesses EET activity that enhances anaerobic growth.^9,10^ We further identified an eight-gene cluster responsible for EET (**Figure 1B**).^11^ Within this cluster, we found that the gene *fmnB* encoded an ApbE-like protein that flavinylated a second protein from the cluster, PplA, at two sites.^11–13^ Homologous genes are present in a number of related Firmicutes and have been implicated in similar EET activities in several bacteria.^14–18^

Extracytosolic proteins acquire cofactors through distinct mechanisms. Some proteins are loaded with their cofactors in the cytosol and then transported by the TAT secretion system across the cytoplasmic membrane in a folded state.^19^ Other proteins are transported across the cytoplasmic membrane in an unfolded state by the Sec secretion system and fold into their active form in the extracytosolic space.^19^ The latter scenario requires an extracytosolic supply of the cofactor, which can be accomplished by transport from the cytosol. For example, the CcsBA transporter transfers heme across the cytoplasmic membrane and is required for loading heme cofactors into cytochromes.^20^

The extracytosolic localization of FmnB and other ApbEs necessitates a source of extracytosolic FAD. We previously proposed that *L. monocytogenes* FAD was supplied by an energy coupling-factor (ECF) transporter that contained FmnA and RibU subunits.^11^ These functional assignments were made because FmnA and RibU exhibit high sequence homology to characterized flavin transporter subunits and because PplA is unflavinylated in Δ*fmnA* and Δ*ribU* strains but rescued by the application of exogenous FAD.^11,21,22^

ECF transporters are a widespread class of bacterial transporters that have been implicated in the import of many metabolites.^23^ ECF transporters are generally comprised of four protein subunits. This includes a pair of ATPases (ECF-A and ECF-A’), a transmembrane domain (ECF-T), and a substrate-binding subunit (ECF-S). While ECF-S subunits that transport different substrates exhibit highly variable sequences, structural studies have revealed a conserved ECF-S fold. ECF transporters within have previously been identified in bacterial genomes on the basis of: (1) gene colocalization (ECF transporter subunits often cluster together in genomes) and (2) the high sequence homology of ECF-T, ECF-A, and ECF-A’ subunits.^23^

Here we reevaluate the basis of FAD secretion in *L. monocytogenes*. We show that RibU is dispensable for FAD secretion when the concentration of cytosolic flavins is normalized and identify a second protein in the EET gene cluster, EetB, that serves as the ECF-S for FAD export. We identify homologous genes in many bacterial genomes and find that they often colocalize with *apbE*, suggesting a conserved role in flavinylation. These studies reveal complex basis of ECF transporter function in flavin acquisition and trafficking.

## Results

### RibU is dispensable for PplA flavinylation when flavin auxotrophy is relieved

Based on the observation that the *L. monocytogenes* Δ*ribU* strain exhibited diminished EET activity and PplA flavinylation, we previously proposed that RibU served as the ECF-S subunit of a putative FAD transporter responsible for the export of FAD from the cytosol to extracytosolic FmnB.^11^ However, we subsequently discovered that RibU transports multiple flavins into the cell and this led us to consider an alternative explanation for the observed phenotypes.^24^ Because *L. monocytogenes* is a flavin auxotroph, impaired flavin import likely leads to a diminished cytosolic flavin level in the Δ*ribU* strain and this could indirectly impair FAD secretion.^24^ To address this possibility, we employed a previously described prototrophic *L. monocytogenes* strain which is relieved of flavin auxotrophy through heterologous expression of the *Bacillus subtilis* riboflavin *ribDEAHT* biosynthesis operon.^24^ Consistent with RibU being important for maintaining cytosolic flavin levels in wildtype *L. monocytogenes*, expressing RibDEAHT in Δ*ribU* restored EET activity and PplA flavinylation to wildtype levels (**Figure 1C & 1D**). By contrast, *ribDEAHT* expression in Δ*fmnB*, which lacks the flavin transferase and is thus not expected to depend upon on cytosolic flavin levels, failed to rescue the defect in EET (**Figure 1C**). These results thus provide evidence that decreased cytosolic flavin concentrations account for the observed Δ*ribU* phenotype and demonstrate that RibU is dispensable for trafficking extracytosolic FAD required for EET.

### FmnA and EetB are required for PplA flavinylation in the absence of exogenous FAD

Having ruled out RibU as a subunit of the FAD exporter, we sought to determine alternative genes essential for FAD secretion. We previously identified *fmnA* as encoding a protein on the EET locus with homology to an ECF-T subunit and found that it was essential for EET activity of cells grown in rich media but that this phenotype could be reversed by the application of exogenous FAD.^11^ We thus asked how the Δ*fmnA* strain responded to engineered flavin prototrophy. In contrast to Δ*ribU*, we found that expressing *ribDEAHT* had no effect on EET activity and PplA flavinylation of the Δ*fmnA* strain (**Figure 1C & 1D**). This result supported our original interpretation of FmnA representing an ECF-T transporter subunit and suggested that its corresponding ECF-S subunit had been previously overlooked.

We next turned to the question of the identity of the ECF-S that acts with FmnA in FAD export. Since the ECF-S subunit of the FAD exporter should be essential for EET activity, we reasoned the ECF-S gene was likely localized to the EET gene cluster. As *eetA* and *eetB* provided the only genes without an assigned function and encode membrane proteins consistent with a transporter function, we reasoned that they presented the strongest candidates. We generated Δ*eetA* and Δ*eetB* strains and tested their EET activity. In contrast to the previously described *eetA* transposon mutant, EET activity of the Δ*eetA* strain did not differ from wildtype, suggesting that the previously observed phenotype may have been due to a polar effect caused by transposon insertion.^11^ By contrast, the Δ*eetB* strain had negligible EET activity and thus continued to present a promising ECF-S candidate (**Figure 1C**).

We next assessed the flavinylation status of PplA in the *ribDEAHT*-expressing Δ*eetB* strain. Proteomic analysis of *L. monocytogenes* cells revealed that PplA was unflavinylated in this strain (**Figure 1D**). Consistent with the lack of PplA flavinylation and impaired EET activity resulting from compromised FAD trafficking, supplementing Δ*eetB* cells with exogenous FAD partially restored EET activity (**Figure 1C**). Importantly, the lack of PplA flavinylation and EET activity of the Δ*eetB* strain was maintained in the RibDEAHT flavin prototroph background, ruling out diminished cytosolic flavin concentrations as the source of the Δ*eetB* phenotype (**Figure 1C**). These findings demonstrate a role for EetB in FAD export.

### EetB is an FAD-binding protein that resembles characterized ECF-Ss

As our genetic studies support the conclusion that *eetB* is essential for FAD secretion, we questioned whether EetB was the FAD-exporting ECF-S. Since experimentally characterized ECF-S subunits with distinct substrate specificities possess low sequence homology but similar tertiary structures, we used the structure modeling software AlphaFold to interrogate structural attributes of EetB.^25–27^ A predicted AlphaFold model of EetB revealed striking structural similarity to experimentally characterized ECF-Ss, including RibU, bolstering the case for the protein possessing an ECF-S functionality (**Figure 2A**).

**Figure 2.**
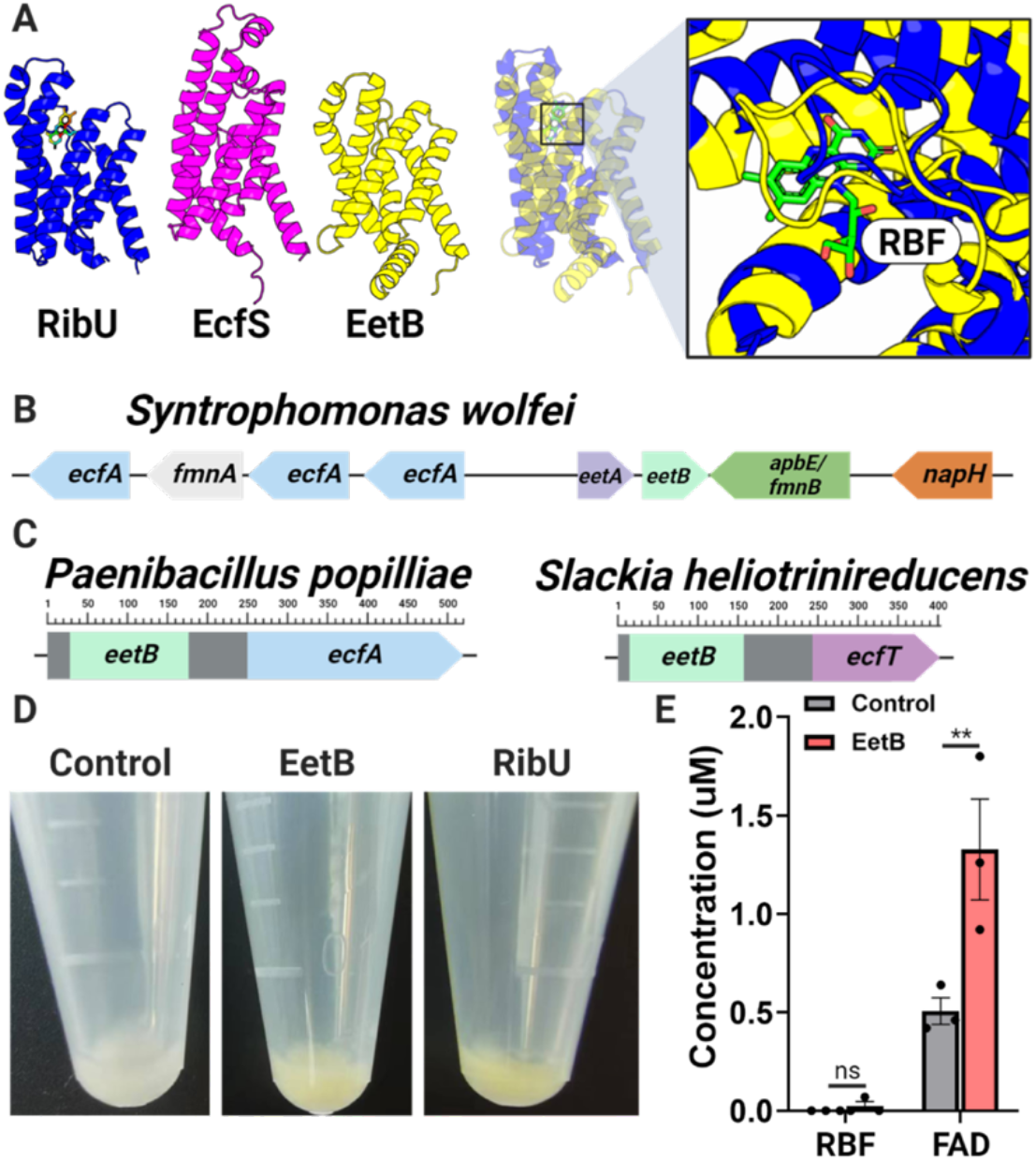
EetB resembles ECF substrate-binding proteins and binds FAD. (A) Comparison of AlphaFold model of EetB to crystal structures of ECF riboflavin and pantothenate substrate-binding subunits, RibU (PDB code 5KBW) and EcfS (PDB code 4RFS), respectively. Riboflavin (RBF) is represented as a stick in the RibU structure. (B) A representative bacterial gene cluster in which *eetB* neighbors genes with ECF-T, ECF-A, and ECF-A’ domains. (C) Domain architecture of representative bacterial proteins (NCBI accessions WP_006284433.1 and WP_012799238.1) that contain *eetB* and ECF domains. (D) Induced *E. coli* cell pellets containing *ribU, eetB*, and plasmid control overexpression vectors. (E) Riboflavin (RBF) and FAD pulled down from *E. coli* cells containing *eetB* and plasmid control overexpression vectors. The means and SEMs of three independent experiments are shown. Statistical significance was determined by performing a *t*-test comparing plasmid control and *eetB* overexpression vectors. **P < 0.01; ns, not significant, P > 0.05.

Genes encoding subunits of characterized ECF transporters are often co-transcribed from an operon. We thus reasoned that the genomic context of *eetB* genes could provide additional evidence of ECF functionality. Indeed, we identified *eetB* genes in several bacterial genomes that colocalized with genes encoding ECF-T and ECF-A/ECF-A’ subunits (**Figure 2B**). Following a similar logic regarding the relationship of functionally related proteins, protein subunits that form a multi-protein complex in one organism are often contained on a single polypeptide chain within other organisms.^28^ The observation of multiple genes from different bacterial genomes that encode proteins with EetB and ECF-T domains (*e*.*g*., NCBI accessions WP_021725833.1, MBE6480012.1, and MBQ3267887.1) or EetB and Ecf-A domains (*e*.*g*., NCBI accessions WP_006284433.1, WP_111154897.1, and WP_143797423.1) thus provides additional support for the attributed role of EetB as an ECF-S (**Figure 2C**).

To address whether EetB might be a flavin-transporting ECF-S similar to RibU, we expressed RibU and EetB in *E. coli*. We observed that both proteins localized to the insoluble fraction of the resulting cell lysates and presented a yellowish hue, consistent with an association with flavins (which are naturally colored yellow) (**Figure 2D**). To address the hypothesized flavin-binding activity of EetB, we measured flavin levels in the insoluble fraction of EetB-overexpressing *E. coli* cells and found that EetB pulled down FAD (**Figure 2E**). Collectively, these analyses demonstrate that EetB is an FAD-binding protein that possesses an ECF-S-like structure consistent with a transport function.

### ECF ATPases are required for PplA flavinylation in the absence of exogenous FAD

Previous ECF transporter characterization has focused on small molecule import. As the putative EetB-FmnA complex is the first transporter identified with apparent export activity, we sought to clarify its mechanism of action. ECF importers typically require two ATPase subunits (ECF-A and ECF-A’) for transporter function. As some bacteria use the same ATPase subunits to engage multiple ECF transport systems, we reasoned that the ATPases that function in the RibU ECF transporter flavin import system might also participate in FAD secretion.^29,30^ To test this hypothesis, we generated an Δ*ecfA*/*ecfA’* strain that lacked both of the previously characterized RibU ATPases.^19^ Consistent with EcfA/EcfA’ being essential for FAD export, the Δ*ecfA*/*ecfA’* strain was deficient for EET activity, PplA flavinylation, and resembled Δ*fmnA* and Δ*eetB* strains in its response to RibDEAHT expression and exogenous FAD application (**Figure 1C**). These findings thus provide evidence that the EcfA/EcfA’ provide dual functions in flavin import and export.

### Structural models illuminate ECF transporters with distinct subunit compositions

Previous studies suggest that RibU, EcfT, EcfA, and EcfA’ form an ECF transporter responsible for riboflavin, FMN, and FAD uptake in *L. monocytogenes*.^22,24,31^ By contrast, the phenotypes for *L. monocytogenes* Δ*eetB*, Δ*fmnA*, Δ*ecfA*/*ecfA’* strains identified in our studies suggest that FAD export occurs through an ECF transporter with EetB, FmnA, EcfA, and EcfA’ subunits (**Figure 1C**). To further address the feasibility of the distinct implied modes of flavin transport, we used the AlphaFold-multimer software to model putative RibU/EcfT/EcfA/EcfA’ and EetB/FmnA/EcfA/EcfA’ complex structures. Both resulting transporter structures exhibit striking similarity to a previously determined crystal structure of the folate ECF transporter (**Figure 3A**). These structural models are thus broadly consistent with the idea that ECF transporters with distinct subunit compositions could be responsible for the observed phenotypes.

**Figure 3.**
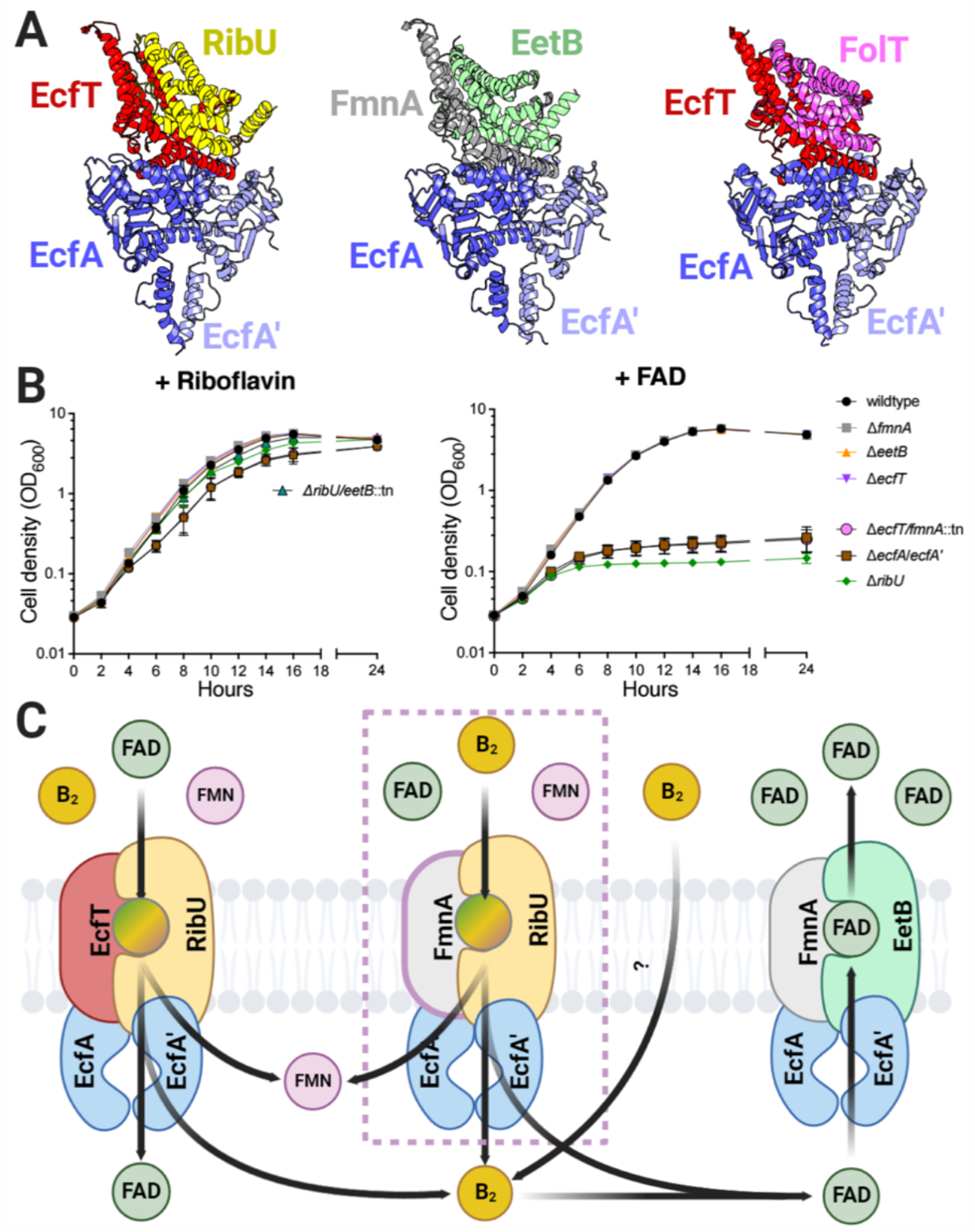
ECF transporter subunits are partially functionally redundant for flavin import and export in *L. monocytogenes*. (A) Comparison of AlphaFold-multimer RibU-EcfT-EcfA-EcfA’ and EetB-FmnA-EcfA-EcfA’ complex models to a crystal structure of the FolT-EcftT-EcfA-EcfA’ folate ECF transporter (PDB code 4HUQ). (B) Growth curves of indicated *L. monocytogenes* strains grown in chemically defined media supplemented with riboflavin as the sole flavin source. Means and SDs from three independent experiments are shown. (C) Growth curves of indicated *L. monocytogenes* strains grown in chemically defined media supplemented with FAD as the sole flavin source. Means and SDs from three independent experiments are shown. (D) Model of ECF complexes that function in *L. monocytogenes* flavin import and export supported observed phenotypes.

### EcfT and FmnA ECF-T subunits are functionally redundant in flavin import

We next sought to address how the direction of transport (import vs. export) could be achieved through related ECF transporters. *L. monocytogenes* is a flavin auxotroph and we previously found that *ribU* was essential for growth in conditions where FMN or FAD are the sole available flavin.^24^ In contrast to these flavin nucleotides, the Δ*ribU* strain lacked a phenotype in the presence of riboflavin.^24^ While it thus seemed plausible that EetB could contribute to riboflavin uptake, we found that a Δ*ribU/eetB*::tn strain grew similarly to wildtype *L. monocytogenes* in chemically defined media that contained riboflavin as the sole flavin (**Figure 3B**). These results suggest that EetB does not substitute for the RibU ECF-S in flavin uptake, and that riboflavin can be imported through a presently unknown alternative mechanism.

We next asked about the functional redundancy of ECF-T subunits. While the observation that FmnA is essential for PplA flavinylation suggests that EcfT cannot substitute in FAD export, we wondered if the converse was true (i.e., whether FmnA could facilitate flavin import in the absence of EcfT). Since RibU was essential for growth when FAD was the sole flavin present, we tested ECF-T mutants in this condition.^24^ Growth of the Δ*ecfT* and Δ*fmnA* strains resembled wildtype *L. monocytogenes* in conditions with riboflavin or FAD (**Figure 3B**). By contrast, the Δ*ecfT/fmnA*::tn strain failed to grow in the presence of FAD, similar to the Δ*ribU* and Δ*ecfA*/*ecfA’* strains (**Figure 3B**). These results suggest that either EcfT or FmnA provides a functionally viable ECF-T for RibU-mediated flavin import and underscore the flexible ECF-T subunit usage for flavin import (**Figure 3C**).

### Gene colocalization suggests a conserved role for EetB in ApbE-associated flavin trafficking

Having established EetB-FmnA function in FAD export, we next sought to address whether this transporter might be important for FAD trafficking in other microbes. We reasoned that the unique substrate-binding subunit provided the clearest marker of likely FAD export activity and thus searched for *eetB* homologs (Pfam accession PF07456) within a collection of 31,910 genomes representative of the genetic diversity of the prokaryotes.^32,33^ We identified 4,214 bacterial genomes that encode an *eetB* homolog. Taxonomic analyses revealed that *eetB* homologs are most abundant in Firmicutes, but also present in a subset of genomes from other phyla (**Figure 4A**).

**Figure 4.**
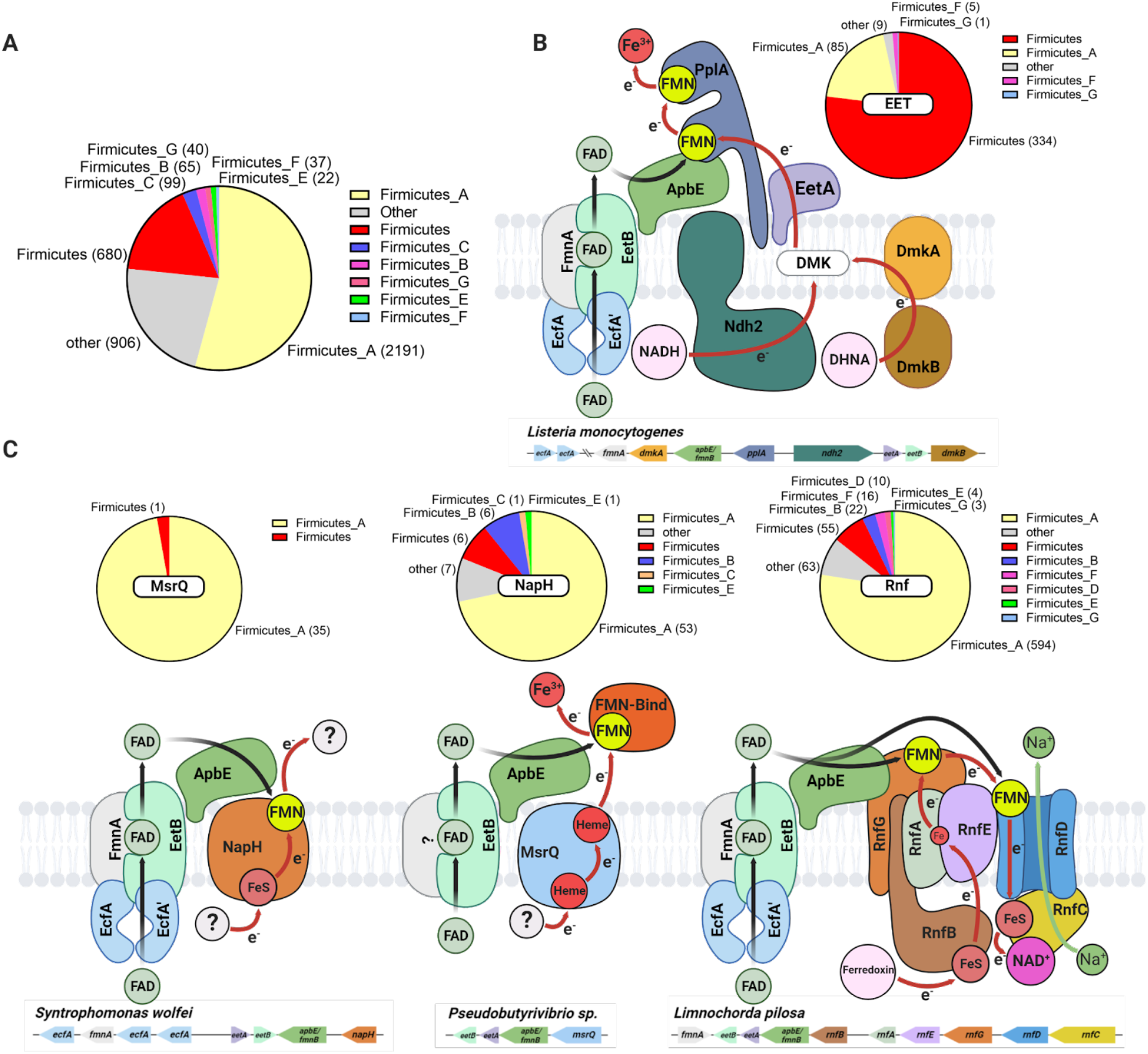
*eetB*s frequently colocalizes with ApbE-associated extracytosolic flavinylation system genes in bacterial genomes. (A) Pie chart showing the number of GTDB reference genomes/phylum that encode an *eetB* homolog. (B) Proposed basis of flavin export for *L. monocytogenes* flavinylation and EET activity. Pie chart shows the number of GTDB reference genomes that contain a gene cluster with both EET and *eetB* genes. (C) Proposed models of flavin export in previously identified flavinylation-associated MsrQ-like, NapH, and Rnf electron transfer systems.^4^ Pie charts show the number of GTDB reference genomes that contain a system gene cluster with an *eetB* homolog.

To clarify the role of EetB in diverse microbes, we next applied a guilt-by-association-based analysis. Our approach took advantage of the frequent colocalization of genes with related functions on prokaryotic genomes and was previously employed to differentiate distinct subtypes of ApbE flavinylation based on colocalization with *apbE*.^3^ Inspecting the genes that colocalized with *eetB*s led to several insights, which are elaborated below.

Supporting the proposed relationship of EetB with other ECF transporter subunits, we found that *eetB* colocalized with ECF-T subunit genes in 291 genomes and ECF-A/ECF-A’ subunits in 796 genomes, often with ECF subunit genes arranged in an apparent operon. Additionally, *eetB* colocalized with *eetA* in 3,328 genomes. While the *L. monocytogenes* Δ*eetA* strain lacked an EET phenotype, this conserved synteny suggests a functional link between EetA and EetB. Further supporting this functional association, we identified genes encoding a single polypeptide chain with both EetA and EetB domains in 14 genomes.

We also found that *eetB* colocalizes with *apbE* on 2,169 genomes. ApbEs have been shown to function in multiple extracytosolic redox activities and we recently proposed operational definitions that enable the identification of gene clusters encoding 10 types of ApbE flavinylated systems with different mechanisms of membrane electron transfer or substrate specificity.^3^ We applied these definitions to determine which types of ApbE flavinylated systems colocalize with *eetB* and found that *eetB* most commonly colocalizes with Rnf and EET systems, but also some MsrQ-like and NapH-like systems (**Figure 4B-4E**). The association with Rnf was particularly pronounced in the class Clostridia, with *eetB* colocalized with Rnf genes in 572 genomes. These results thus suggest that ECF exporters traffic FAD to a functionally diverse subset of bacterial ApbEs.

## Discussion

The studies presented here establish the basis of FAD trafficking in *L. monocytogenes* and, notably, provide the most extensive evidence of the role of ECF-like transporters in small molecule export. Strikingly, our comparison of strains deficient in various ECF components supports the existence of distinct import and export transporters that share subunits (ECF-AA’) and exhibit partial functional redundancies (import ECF-Ts). These findings thus suggest a complex basis of bidirectional flavin transport across the *L. monocytogenes* cytoplasmic membrane (**Figure 3D**).

While little is known about the mechanism of ECF export, previous research has generated considerable evidence about the basis of ECF import. These studies reveal that ECF-S remains monomeric in the absence of substrate, likely adopting an ‘outward’ facing orientation that enables substrate binding from the extracytosolic space.^22,34,35^ Once substrate is bound, ECF-S undergoes a ‘toppling’ conformational change to an ‘inward’ orientation where it engages the ECF-T, ECF-A, ECF-A’ complex in a manner that facilitates substrate release into the cytosol. ATP hydrolysis in the ECF complex then causes release of apo ECF-S, which reverts to its monomeric ‘outward’ orientation (**Figure 5A**).^22,35^ Considering the similarity of subunits, the proposed mechanism of ECF import has implications for how EetB might function in FAD export. Simply reversing the relationship between ECF-S with the rest of the ECF transporter complex could reverse the direction of transport (**Figure 5B**). While additional studies will be necessary to definitively address transport mechanisms, ECF export may differ from ECF import in inverting the ECF complex’s unliganded versus substrate-bound ECF-S affinity and the kinetics of ATP hydrolysis/ECF-S release.

**Figure 5.**
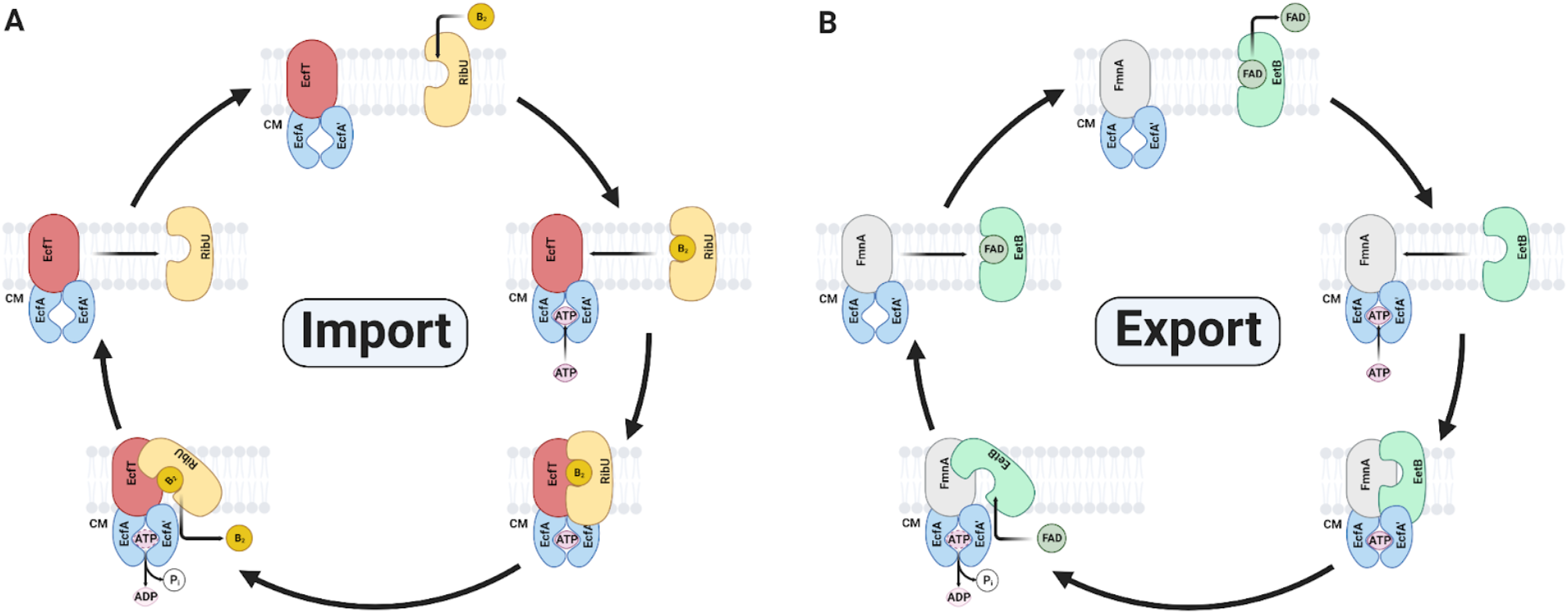
Proposed basis of bidirectional ECF transport. (A) Summary of previously proposed models of ECF import. (B) Speculative model of ECF export illustrating how directionality of transport could be achieved through a broadly conserved mechanism.

An interesting aspect of *eetB* and other flavin exporter genes identified in our bioinformatic analyses regards their frequent colocalization with an *apbE* gene. This distinguishes the ECF transporter from a previously identified flavin exporter that similarly traffics flavins in some Gram-negative bacteria.^2,3^ The *eetB*-*apbE* association suggests that EetB may be adapted for efficient flavin delivery to ApbE. Indeed, a regulatory mechanism that enabled targeted delivery to ApbE might explain why, despite several efforts, we were unable to detect differences in the level of extracellular flavins within *eetB*-deficient strains. Flavin delivery to ApbE may thus provide an attractive model for future investigations into the mechanism of targeted extracytosolic cofactor delivery in bacteria.

## Methods

### Bacterial strains and culture

All strains of *L. monocytogenes* used in this study (**Table S1**) were derived from the wildtype 10403S strain and cultured in a previously described chemically defined synthetic media containing 200 µg/mL of streptomycin.^36,37^ Deletion of genes was done using allelic exchange with the temperature-sensitive plasmid pKSV7, as previously described.^38^ Generation of the strains expressing the *ribDEAHT* operon from the constitutive promoter pHyper (pHyper *ribDEAHT* construct) was done by amplifying the *ribDEAHT* operon from *B. subtilis* and cloning it into the site-specific pPL2 integrating vector. The plasmids were then introduced into *L. monocytogenes* by conjugation, as previously described.^39^ Broth growth curves were performed with *L. monocytogenes* strains from overnight cultures grown in chemically defined synthetic media at 37 °C with shaking (200 rpm). Growth was measured by optical density at a wavelength of 600 nm (OD_600_) and the growth curves were started at an OD_600_ of 0.03.

### Ferric iron reductase activity assays

Strains were grown to mid-log phase (OD_600_ = ~0.4-0.6) in chemically defined synthetic media, or chemically defined synthetic media supplemented with 0.5 mM FAD, washed twice, then normalized to an OD_600_ of 0.5. One mL of the washed bacteria was then resuspended in 4 mM Ferrozine in chemically defined synthetic media. To conduct the assay, 100 μL of resuspended bacteria were mixed with 100 μL of 100 mM ferric ammonium citrate, in chemically defined synthetic media, and transferred into a 96-well plate format in triplicate. Measurements were done using a plate reader with the temperature set at 37°C and absorbance was read at 560 nm every 30 seconds for 1.5 hours. Maximal rates (typically over 2 min) were calculated and reported as the percent ferric iron reductase activity of wildtype.

### Recombinant *ribU* and *eetB* expression and FAD pulldown

*L. monocytogenes* 10403S *eetB* and *ribU* genes were cloned into the pMCSG53 vector. Resulting constructs were transformed into *E. coli* BL21 cells. Overnight cultures of *E. coli* BL21 containing pMCSG53::*empty*, pMCSG53::*eetB*, and pMCSG53::*ribU* were diluted in 5 mL of Luria-Bertani broth (LB) with a final OD_600_ of 0.05. When cell growth reached log phase, FAD was added to a final concentration of 1 μM and protein expression was induced with isopropyl β-D-1-thiogalactopyranoside (IPTG) at a final concentration of 1 mM. Induced cultures were grown overnight at 30°C. Cultures were normalized to an OD_600_ of 1.0 and then 1 mL of cells was collected by centrifugation at 22,100 x g for 1 min. Cell pellets were washed twice with 1 mL of diH_2_O and resuspended in 190 μL of diH_2_O to remove free FAD. To facilitate the release of protein-bound FAD, cell suspensions were incubated at 100°C for 20 minutes and centrifuged at 22,100 x g for 1 minute. Supernatants were collected for the analysis of flavin content.

### Liquid chromatography-mass spectrometry for the detection of flavins

Samples were incubated at -80 °C for at least one hour or up to overnight. Extraction solvent (4 volumes of 100% methanol spiked with internal standards and stored at -80 °C) was added to the liquid sample (1 volume) in a microcentrifuge tube. Tubes were then centrifuged at -10 °C, 20,000 x g for 15 min, and supernatant was used for subsequent metabolomic analysis.

Samples were dried down completely using a Genevac EZ-2 Elite. Samples were resuspended in 50 µL of 50:50 water:methanol and added to an Eppendorf thermomixer® at 4 °C, 1000 rpm for 15 min to resuspend analytes. Samples were then centrifuged at 4 °C, 20,000 x g for 15 min to remove insoluble debris, and 40 µL of supernatant was transferred to a 96 deep-well plate (Agilent 5065-4402). Samples were analyzed on an Agilent 1290 infinity II liquid chromatography system coupled to an Agilent 6470 triple quadrupole mass spectrometer, operating in positive mode, equipped with an Agilent Jet Stream Electrospray Ionization source (LC-ESI-QQQ). Each sample (2 µL) was injected into a Acquity UPLC HSS PFP column, 1.8 µm, 2.1 × 100 mm (Waters; 186005967), equipped with a Acquity UPLC HSS PFP VanGuard Pre-column, 100Å, 1.8 μm, 2.1 mm × 5 mm (Waters; 186005974), at 45 °C. Mobile phase A was 0.35% formic acid in water and mobile phase B was 0.35% formic acid in 95:5 acetonitrile:water. The flow rate was set to 0.5 mL/min starting at 0% B held constant for 3 min, then linearly increased to 50% over 5 min, then linearly increased to 95% B over 1 min, and held at 100% B for the next 3 min. Mobile phase B was then brought back down to 0% over 0.5 min and held at 0% for re-equilibration for 2.5 min. The QQQ electrospray conditions were set with capillary voltage at 4 kV, nozzle voltage at 500 V, and Dynamic MRM was used with a cycle time of 500 ms. Transitions were monitored in positive mode for two analytes, riboflavin and flavin adenine dinucleotide (FAD). The transitions for riboflavin and FAD were 377.1 m/z to 243 m/z and 786.1 m/z to 348 m/z, respectively. Authentic standards were purchased for riboflavin (Supelco; riboflavin B2) and FAD (Sigma-Aldrich; flavin adenine dinucleotide disodium salt hydrate) to make 1 mg/mL-stock solutions in methanol. These solutions were used to prepare a 10-point calibration curve, ranging from 3.1 nM to 0.2 mM for riboflavin and 3.9 nM to 1 mM for FAD. Data analysis was performed using MassHunter Quant software (version B.10, Agilent Technologies) and confirmed by comparison with standards. Normalized peak areas were calculated by dividing raw peak areas of the targeted analytes by averaged raw peak areas of two internal standards (melatonin and kynurenic acid).

### Transporter structural models

The AlphaFold model of EetB (accession AF-Q927J9-F1) was downloaded from the Uniprot database.^40^ RibU-EcfT-EcfA-EcfA’ and EetB-FmnA-EcfA-EcfA’ complex models were generated with AlphaFold-multimer software using default settings in the ColabFold platform.^41,42^

### Liquid chromatography-mass spectrometry analysis of trypsin-digested proteins

Samples of trypsin-digested proteins were analyzed using a Synapt G2-Si ion mobility mass spectrometer that was equipped with a nanoelectrospray ionization source (Waters, Milford, MA). The mass spectrometer was connected in line with an Acquity M-class ultra-performance liquid chromatography system that was equipped with trapping (Symmetry C18, inner diameter: 180 μm, length: 20 mm, particle size: 5 μm) and analytical (HSS T3, inner diameter: 75 μm, length: 250 mm, particle size: 1.8 μm) columns (Waters). Data-independent, ion mobility-enabled, high-definition mass spectra and tandem mass spectra were acquired in the positive ion mode.^43–46^ Data acquisition was controlled using MassLynx software (version 4.1) and tryptic peptide identification and relative quantification using a label-free approach were performed using Progenesis QI for Proteomics software (version 4.0, Waters).^47–49^ Data were searched against the *Listeria monocytogenes* serotype 1/2a (strain 10403S) protein database to identify tryptic peptides, with carbamidomethylcysteine as a fixed post-translational modification and methionine sulfoxide and threonine flavinylation as variable post-translational modifications.^50^ Calculation of the percentage of flavinylation for each bacterial strain was performed by dividing the abundance of a residue/peptide bearing a flavinylation modification by the total abundance and multiplying by 100.

### Genome collection for analysis of *eetB* gene clusters

The 30,238 bacterial and 1672 archaeal genomes from the GTDB (release 05-RS95 of July 17, 2020) were downloaded with the taxonomy and predicted protein sequences.^32,33^

### Functional annotation of *eetB* gene clusters

Protein sequences were functionally annotated based on the accession of their best Hmmsearch match, version 3.3 (E-value cut-off 0.001)^51^ against the KOfam database (downloaded on February 18, 2020).^52^ Domains were predicted using the same Hmmsearch procedure against the Pfam database, version 33.0.^53^ SIGNALP, version 5.0, was run to predict the putative cellular localization of the proteins using the parameters -org arch in archaeal genomes and -org gram+ in bacterial genomes.^54^ Prediction of transmembrane helices in proteins was performed using TMHMM, version 2.0 (default parameters).^55^

### Detection of *eetB* gene clusters and association with flavinylated systems

The five genes downstream and upstream of an *eetB* (Pfam accession PF07456) encoding genes were first collected. The *eetB* gene clusters were then assigned to flavinylated systems based on the presence of previously reported key genes.^4^

## Acknowledgments

Research reported in this publication was supported by funding from the National Institutes of Health (1P01AI063302 & 1R01AI27655 to D.A.P., K22AI144031 to S.H.L, and 1S10OD020062-01 to A.T.I.), the National Academies of Sciences, Engineering, and Medicine (Ford Foundation Fellowship to R.R.-L.), the University of California Dissertation-Year Fellowship (to R.R.-L.), and the Searle Scholars Program (to S.H.L). We thank the Host-Microbe Metabolomics Facility for experimental assistance.

